# Connecting actin polymer dynamics across multiple scales

**DOI:** 10.1101/2020.01.28.923698

**Authors:** Calina Copos, Brittany Bannish, Kelsey Gasior, Rebecca L. Pinals, Minghao W. Rostami, Adriana Dawes

## Abstract

Actin is an intracellular protein that constitutes a primary component of the cellular cytoskeleton and is accordingly crucial for various cell functions. Actin assembles into semi-flexible filaments that cross-link to form higher order structures within the cytoskeleton. In turn, the actin cytoskeketon regulates cell shape, and participates in cell migration and division. A variety of theoretical models have been proposed to investigate actin dynamics across distinct scales, from the stochastic nature of protein and molecular motor dynamics to the deterministic macroscopic behavior of the cytoskeleton. Yet, the relationship between molecular-level actin processes and cellular-level actin network behavior remains understudied, where prior models do not holistically bridge the two scales together.

In this work, we focus on the dynamics of the formation of a branched actin structure as observed at the leading edge of motile eukaryotic cells. We construct a minimal agent-based model for the microscale branching actin dynamics, and a deterministic partial differential equation model for the macroscopic network growth and bulk diffusion. The microscale model is stochastic, as its dynamics are based on molecular level effects. The effective diffusion constant and reaction rates of the deterministic model are calculated from averaged simulations of the microscale model, using the mean displacement of the network front and characteristics of the actin network density. With this method, we design concrete metrics that connect phenomenological parameters in the reaction-diffusion system to the biochemical molecular rates typically measured experimentally. A parameter sensitivity analysis in the stochastic agent-based model shows that the effective diffusion and growth constants vary with branching parameters in a complementary way to ensure that the outward speed of the network remains fixed. These results suggest that perturbations to microscale rates can have significant consequences at the macroscopic level, and these should be taken into account when proposing continuum models of actin network dynamics.

## 1 Introduction

A cell’s mechanical properties are determined by the cytoskeleton whose primary components are actin filaments (F-actin) [1–4]. Actin filaments are linear polymers of the abundant intracellular protein actin [5–7], referred to as G-actin when not polymerized. Regulatory proteins and molecular motors constantly remodel the actin filaments and their dynamics have been studied *in vivo* [8], in reconstituted *in vitro* systems [2, 9], and *in silico* [10]. Actin filaments are capable of forming large-scale networks and can generate pushing, pulling, and resistive forces necessary for various cellular functions such as cell motility, mechanosensation, and tissue morphogenesis [8]. Therefore, insights into actin dynamics will advance our understanding of cellular physiology and associated pathological conditions [2, 11].

Actin filaments in cells are dynamic and strongly out of equilibrium. The filaments are semi-flexible, rod-like structures approximately 7 nm in diameter and extending several microns in length, formed through the assembly of G-actin subunits [6, 7]. A filament has two ends, a barbed end and a pointed end, with distinct growth and decay properties. A filament length undergoes cycles of growth and decay fueled by an input of chemical energy, in the form of ATP, to bind and unbind actin monomers [6, 7]. The rates at which actin molecules bind and unbind from a filament have been measured experimentally [3, 12, 13]. The cell tightly regulates the number, density, length, and geometry of actin filaments [7]. In particular, the geometry of actin networks is controlled by a class of accessory proteins that bind to the filaments or their subunits. Through such interactions, accessory proteins are able to determine the assembly sites for new filaments, change the binding and unbinding rates, regulate the partitioning of polymer proteins between filaments and subunit forms, link filaments to one another, and generate mechanical forces [6, 7]. Actin filaments have been observed to organize into branched networks [14, 15], sliding bundles extending over long distances [16], and transient patterns including vortices and asters [17, 18].

To generate pushing forces for motility, the cell uses the energy of growth or polymerization of Factin [19, 20]. The directionality of pushing forces produced by actin polymerization originates from the uniform orientation of polymerizing actin filaments with their barbed ends towards the leading edge of the cell [8]. Here, cells exploit the polarity of filaments, since growth dynamics are much faster at barbed ends than at pointed ends [21, 22]. Polymerization of individual actin filaments produces piconewton forces [23], and filaments are organized into parallel bundles in filopodia or a branched network in lamellipodia [15]. Lamellipodia are flat cellular protrusions found at the leading edge of motile cells and serve as the major cellular engine to propel the leading edge forward forward [8,24]. Microscopy of lamellipodial cytoskeleton has revealed multiple branched actin filaments [15]. The branching structure is governed by the Arp2/3 protein complex, which binds to an existing actin filament and initiates growth of a new “daughter” filament through a nucleation site at the side of preexisting filaments. Growth of the “daughter” filament occurs at a tightly regulated angle of 70° from the “parent” filament due to the crystal structure of the Arp2/3 complex [25]. The localized kinetics of growth, decay, and branching of a protrusive actin network provide the cell with the scaffold and the mechanical work needed for directed movement.

Many mathematical models have been developed to capture the structural formation and force generation of actin networks [20,26,27]. Due to the multiscale nature of actin dynamics, two main approaches are used: agent-based methods [27–29] and deterministic models using partial differential equations (PDEs) [30–33]. While both techniques are useful for understanding actin dynamics, each presents limitations. Agent-based models more closely capture the molecular dynamics of actin by explicitly considering the behavior of actin molecules through rules, such as, bind to the closest filament at a particular rate. In general, agent-based models simulate the spatiotemporal actions of certain microscopic entities, or “agents”, in an effort to recreate and predict more complex large-scale behavior. In these simulations, agents behave autonomously and through simple rules prescribed at each time step. The technique is stochastic and can be interpreted as a coarsening of Brownian and Langevin dynamical models [34]. However, agent-based approaches are computationally expensive: at every time step, they specifically account for the movement and interaction of individual molecules, while also assessing the effects of spatial and environmental properties that ultimately result in the emergence of certain large-scale phenomena, such as crowding. Such approaches benefit from the direct relationship to experimental measurements of parameters, yet they present a further computational cost in that many instances of a simulation are needed for reliable statistical information. Agent or rule-based approaches have been used to reveal small-scale polymerization dynamics in actin polymer networks [26, 35], but due to the inherent computational complexity, it remains unclear how this information translates to higher length scales, such as the cell, tissue, or whole organism.

To overcome such computational costs and still gain a mechanistic understanding of actin processes, one approach is to write deterministic equations that “summarize” all detailed stochastic events. These approaches rely on differential equations to predict a coarse-grained biological behavior by assuming a well-mixed system where the molecules of interest exist in high numbers [20, 36–38]. In continuum models, the stochastic behavior of the underlying molecules are typically ignored. The challenge lies in determining the terms and parameters of these equations that are representative of the underlying physical system. Thus, these methods use phenomenological parameters of the actin network, such as bulk diffusion and reaction terms, that are less readily obtained experimentally. The relationship between molecular-level actin processes and cellular-level actin network behavior remains disconnected. This disconnect presents a unique challenge in modeling actin polymers in an active system across length scales.

In this work, we design a systematic and rigorous methodology to compare and connect actin molecular effects in agent-based stochastic simulations to macroscopic behavior in deterministic continuum equations. Measures from these distinct-scale models enable extrapolation from the molecular to the macroscopic scale by relating local actin dynamics to phenomenological bulk parameters. First, we characterize the dynamics of a protrusive actin network in free space using a minimal agent-based model for the branching of actin filaments from a single nucleation site based on experimentally measured kinetic rates. Second, in the macroscopic approach, we simulate the spread of actin filament concentration from a source point using a PDE model. The model equation is derived from first principles of actin filament dynamics and is found to be Skellam’s equation for unbounded growth of a species together with spatial diffusion. To compare the emergent networks, multiple instances of the agent-based approach are simulated, and averaged effective diffusion coefficient and reaction rate are extracted from the mean displacement of the advancing network front and from the maximum gradient of the averaged network density. We identify two concrete metrics, mean displacement and maximum gradient of the averaged filament length density, that connect phenomenological bulk parameters in the reaction-diffusion systems to the molecular biochemical rates of actin binding, unbinding, and branching. Using sensitivity analysis on these measures, we demonstrate that the outward movement of the actin network is insensitive to changes in parameters associated with branching, while the bulk growth rate and diffusion coefficient do vary with changes in branching dynamics. We further find that the outward speed, growth rate constant, and effective diffusion increase with F-actin polymerization rate but decrease with increasing depolymerization of actin filaments. By formalizing the relationship between micro- and macro-scale actin network dynamics, we demonstrate a nonlinear dependence of bulk parameters on molecular characteristics, indicating the need for careful model construction and justification when modeling the dynamics of actin networks.

## 2 Mathematical Models

### 2.1 Microscale Agent-Based Model

#### 2.1.1 Model Description

We build a minimal, agent-based model sufficient to capture the local microstructure of a branching actin network. This model includes the dynamics of actin filament polymerization, depolymerization, and branching from a nucleation site [1, 2, 39, 40]. We treat F-actin filaments as rigid rods. Each actin filament has a base (pointed end) fixed in space and a tip (barbed end) capable of growing or shrinking due to the addition or removal of actin monomers, respectively. We assume that there is an unlimited pool of actin monomers available for filament growth, in line with normal, intracellular conditions [3]. For simplicity, we neglect effects of barbed end capping, mechanical response of actin filaments, resistance of the plasma membrane, cytosolic flow, and molecular motors and regulatory proteins. The physical setup of the model is similar to conditions associated with *in vitro* experiments, as well as initial actin network growth in cells, before components such as actin monomers become limiting.

#### 2.1.2 Numerical Implementation

At the start of each simulation, an actin filament of length zero is randomly assigned an angle of growth from the nucleation site located at the origin. Once a filament is prescribed a direction of growth, it will not change throughout the time-evolution of that particular filament. At each subsequent time step in the simulation, there are four possible outcomes: (i) growth of the filament with probability *p*_poly_, (ii) shrinking of the filament with probability *p*_depoly_, (iii) no change in filament length, or (iv) branching of a preexisting filament into a “daughter” filament with probability *p*_branch_ provided that the “parent” filament has reached a critical length *L*_branch_, measured from the closest branch point. To determine which outcome occurs, two random numbers are independently generated for each filament. The first random number governs polymerization (i) or depolymerization (ii): if the random number is less than *p*_poly_, polymerization occurs, and if greater than 1 − *p*_depoly_, depolymerization occurs. If the first random number is greater than or equal to *p*_poly_ and less than or equal to 1 −*p*_depoly_, then the filament neither polymerizes nor depolymerizes this time step, and therefore remains the same length (iii). Both filament growth and shrinking occur in discrete increments corresponding to the length of a G-actin monomer, ∆*x* = 2.7 nm [4]. We enforce that a filament of length zero cannot depolymerize.

The second random number pertains to filament branching (iv). For filaments of length greater than *L*_branch_, a new filament can be initiated at a randomly oriented 70° -angle from a preexisting filament tip in correspondence with the effect of Arp2/3 protein complex. If the second random number is less than the probability of branching, *p*_branch_, for the given filament, then the filament will branch and create a “daughter” filament now capable of autonomous growth and branching. This branching potential models the biological effect of the Arp2/3 complex without explicitly including Arp2/3 concentration as a variable.

This step-wise process is repeated until the final simulation time is reached. Simulation steps are summarized graphically in Fig. 1. All parameters for the model are listed in Table 1. We calculate several different measurements from the microscale simulation, as described below.

**Table 1:** Microscale model parameter values. Details on parameter estimation are available in Section 2.1.3. Values flagged with one star (^*∗*^) were calculated from [3]. For the value flagged with a dagger (^†^): literature measurements of actin filament length per branch vary from 0.02–5 *µ*m [15, 41–45].

| Parameter | Meaning | Value |
| --- | --- | --- |
| $p_{\text{branch}}$ | Probability of branching | normal CDF |
| $p_{\text{poly}}$ | Probability of polymerizing | 0.6* |
| $p_{\text{depoly}}$ | Probability of depolymerizing | 0.02* |
| $L_{\text{branch}}$ | Critical length before branching can occur | $0.2 \mu\text{m}^\dagger$ |
| $T_{\text{end}}$ | Total run time | 10 s |
| $\Delta t$ | Time step | 0.005 s |
| $n_{\text{sim}}$ | Number of independent simulations | 10 |

**Figure 1:**
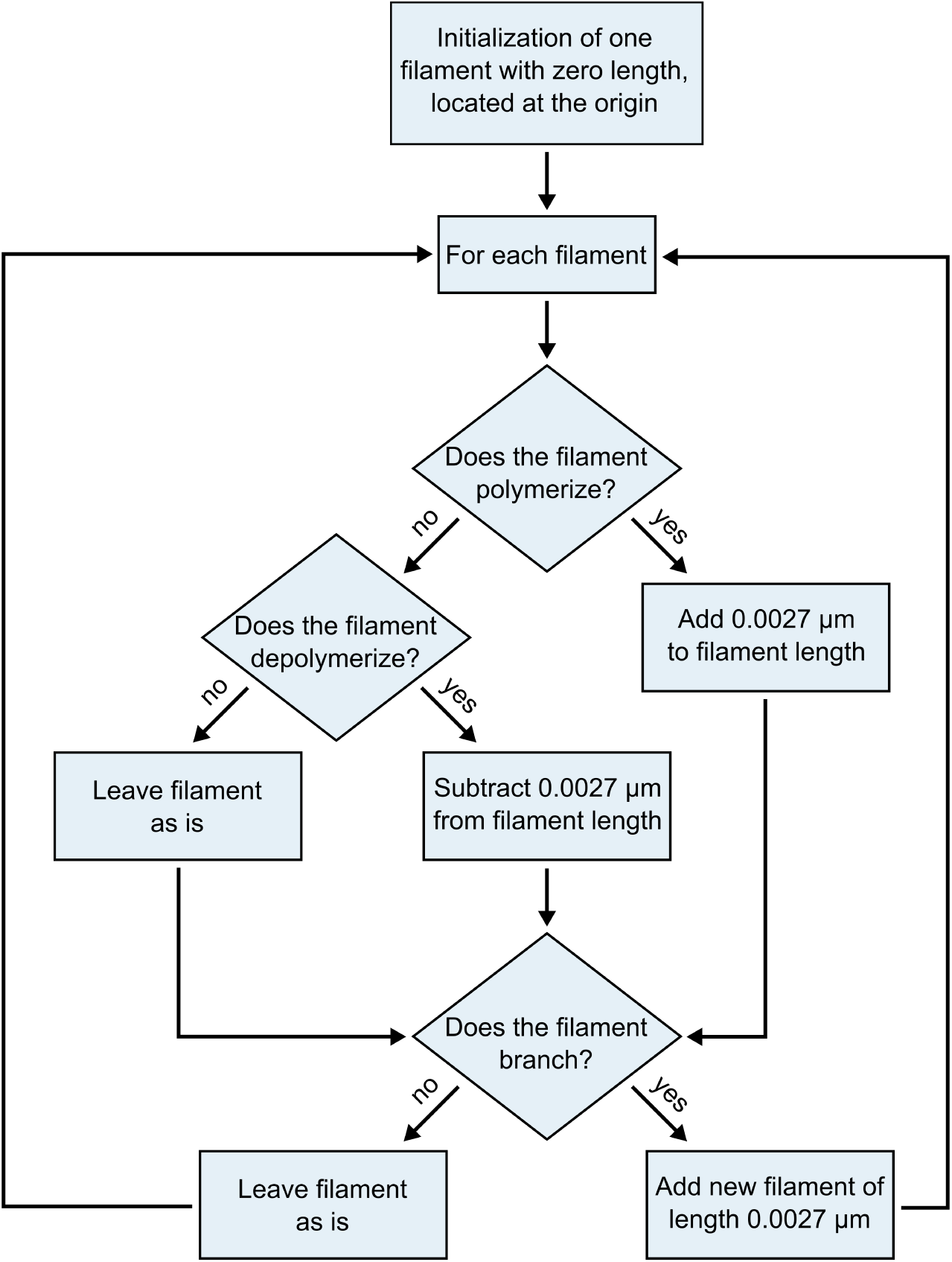
Flow chart of the algorithm implemented for the agent-based microscale model. All steps following “Initialization” are repeated at every time step.

#### 2.1.3 Parameter Estimation

Actin dynamics have been extensively studied *in vivo* and *in vitro*, identifying many rate constants used in this study. Actin monomer elongate the barbed ends of F-actin filaments at a reported velocity of 0.3 *µ*m/s [3]. We use this measurement to calculate the polymerization probability, *p*_poly_, via the formula:

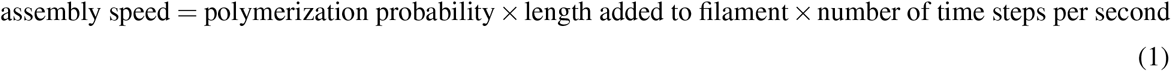

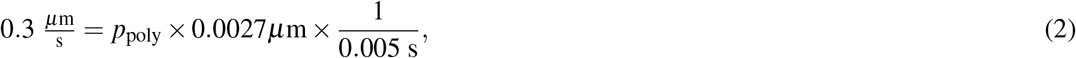

which implies that *p*_poly_ = 0.56. For simplicity, we round this probability to *p*_poly_ = 0.6 in the microscale model simulations. ADP-actin has a depolymerization rate of 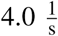 [3]. This value represents the rate of depolymerization of one actin subunit per second, thus a filament loses length at a velocity of:

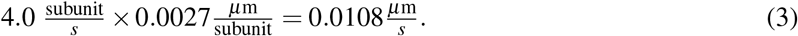

To calculate the depolymerization probability, *p*_depoly_, we use the analogous formula:

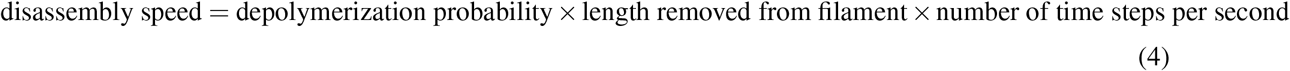

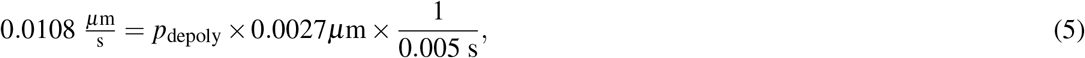

which yields *p*_depoly_ = 0.02.

Model parameter *L*_branch_ represents the critical length a filament must reach before branching can occur. Literature estimates for the spacing of branching Arp2/3 complexes along a filament vary widely, from 0.02 *µ*m to 5 *µ*m [15, 41–45]. We choose an intermediate estimate, *L*_branch_ = 0.2 *µ*m, which is of similar order to the values from other studies [44, 45]. The branching probability, *p*_branch_, is chosen from a normal cumulative distribution function (CDF) with mean, *µ* = 2 and standard deviation, *σ* = 1. For *in vitro* systems, branch formation is inefficient because the reported branching rate once an Arp2/3 complex is bound to a filament is slow (estimated to be 0.0022 *−* 0.007 s^*−*1^) [45]. Given the relative dynamic scales of polymerization/depolymerization versus branching, we assume that (de)polymerization occurs at a prescribed rate, but because branching is infrequent, its probability is drawn from a distribution function.

### 2.2 Macroscale Deterministic Model

#### 2.2.1 Model Description

We model the growth and spread of a branching actin network through a reaction-diffusion form of the chemical species conservation equation derived in Section 2.2.2. This form is frequently known as Skellam’s equation, applied to describe populations that grow exponentially and disperse randomly [46]. Skellam’s equation in 2D is given as follows:

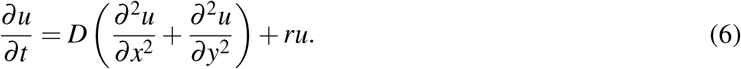

In the context of our actin network, *u*(*x, y,t*) is the density of polymerized actin filaments at location (*x, y*) and time *t*, *D* is the diffusion coefficient of the network (network spread), and *r* is the effective growth rate constant (network growth). Note that the diffusion coefficient is in reference to the bulk F-actin network spread, rather than representing Fickian behavior of monomers as has been done in previous literature [31, 38]. We use no flux boundary conditions in Eq. 6 to enforce no flow of actin across the cell membrane.

#### 2.2.2 Derivation of Reaction Term from First Principles

We present a derivation of the reaction term in Skellam’s continuum description (Eq. 6) from simple kinetic considerations of actin filaments which include polymerization, depolymerization, and branching.

First, we write the molecular scheme for actin filament polymerization and depolymerization in the form of chemical equations. We denote a G-actin monomer in the cytoplasmic pool by *M*, an actin polymer chain consisting of *n* − 1 subunits by *P*_*n*−1_, and a one monomer longer actin polymer chain by *P*_*n*_. The process of binding and unbinding of an actin monomer is described by the following reversible chemical reaction:

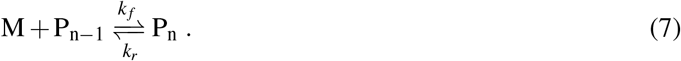

The constants *k*_*f*_ and *k*_*r*_ represent the forward and reverse rate constants, respectively, and encompass the dynamics that lead to the growth/shrinking of an actin filament.

Second, the biochemical reaction in Eq. 7 can be translated into a differential equation that describes rates of change of the F-actin network concentration. To write the corresponding equations, we first use the law of mass action which states that the rate of reaction is proportional to the product of the concentration. Then, the rates of the forward (*r*_*f*_) and reverse (*r*_*r*_) reactions are:

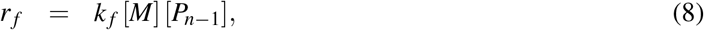

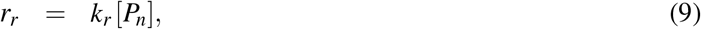

where brackets denote concentrations of *M*, *P*_*n*−1_, and *P*_*n*_. Under the assumption that the forward and reverse reactions are each elementary steps, the net reaction rate is:

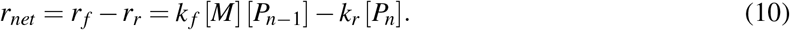

We note that the monomer concentration [*M*] can be eliminated from Eq. 10 if it is expressed in terms of initial concentration of monomers in the cell cytoplasm [*M*_0_]:

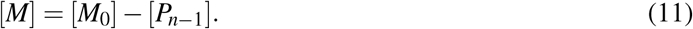

Lastly, we set [*P*_*n*−1_] = [*P*_*n*_] because concentrations of different polymerized actin filaments are chemically indistinguishable for the purpose of this derivation. Then, Eq. 10 becomes

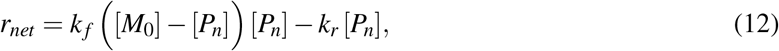

and can be further simplified if we divide both sides of the equation by [*M*_0_]:

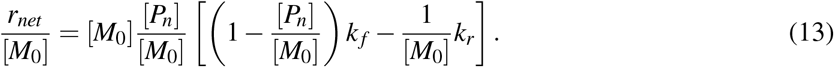

We introduce variable *u* as the polymerized actin concentration normalized by the initial monomeric actin concentration and *r* as an effective growth rate constant:

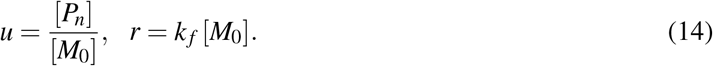

The normalized net reaction rate in Eq. 13 simplifies to

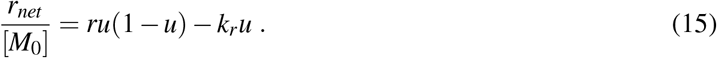

Describing the macroscale dynamics of polymerized actin concentration [*P*_*n*_] as simultaneously diffusing in two-dimensional space with diffusion coefficient *D* and undergoing molecular reactions with the net reaction rate *r*_*net*_ results in the following partial differential equation:

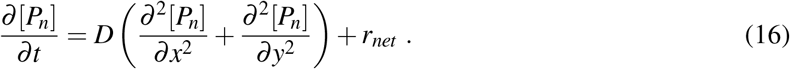

The equation expressed in terms of variable *u* = [*P*_*n*_]/[*M*_0_] becomes:

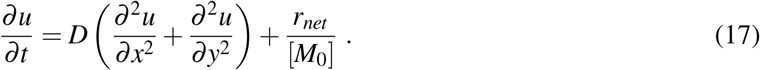

Finally, substituting in the full form of the normalized net reaction rate from Eq. 15 yields the following partial differential equation:

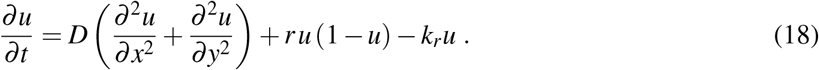

##### Special Cases

We consider two special cases of these kinetics. In the case of normal intracellular conditions [3] where monomeric actin concentration is much larger than polymerized actin concentration, the normalized net reaction rate in Eq. 15 becomes

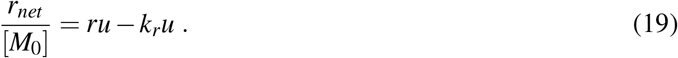

Similarly, in the case of a slow reversal reaction (i.e, slow depolymerization rate with *k*_*r*_ → 0), the normalized net reaction rate in Eq. 15 simplifies to

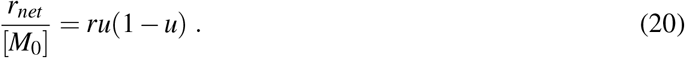

Taking these two special cases together yields the following net reaction rate:

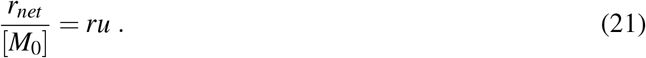

Substituting *r*_*net*_/[*M*_0_] for the case of both unlimited monomers and slow depolymerization into Eq. 17 yields the same functional form as Skellam’s equation for unbounded growth (Eq. 6):

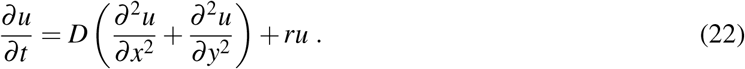

Conversely, substituting *r*_*net*_/[*M*_0_] for only the case of slow depolymerization into Eq. 17 produces Fisher’s equation for saturated growth:

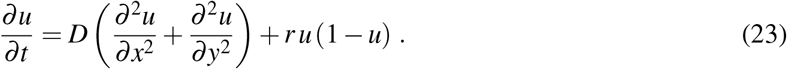

For the current system under study, the two aforementioned special case assumptions hold, in that the monomer pool is unlimited and the rate of polymerization far exceeds the rate of depolymerization. Therefore, the former equation (Skellam’s) is chosen to model macroscale actin dynamics.

#### 2.2.3 Analytical Solution of PDE

The analytic solution to Eq. 6 is:

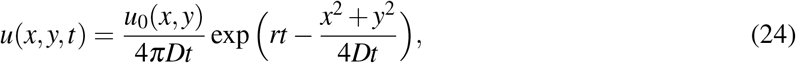

where *u*_0_(*x, y*) is the initial density at (*x, y*). An interesting result of Eq. 24 is that for sufficiently large times *t*, F-actin density *u*(*x, y,t*) propagates as a unidirectional wave moving at a constant speed *v*. To see this, we first fix *u* and solve for *x*^2^ + *y*^2^ from Eq. (24):

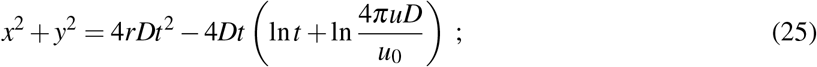

then, we compute the following limit:

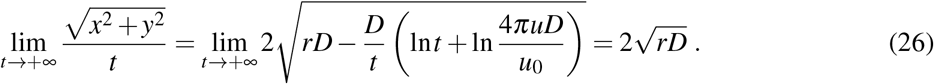

For large enough *t* values, *u*(*x, y,t*) is a traveling wave propagating with speed 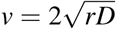. Indeed, in the stochastic simulations, we observe that the speed at which the periphery of the actin network advances is roughly constant (see Fig. 2c).

**Figure 2:**
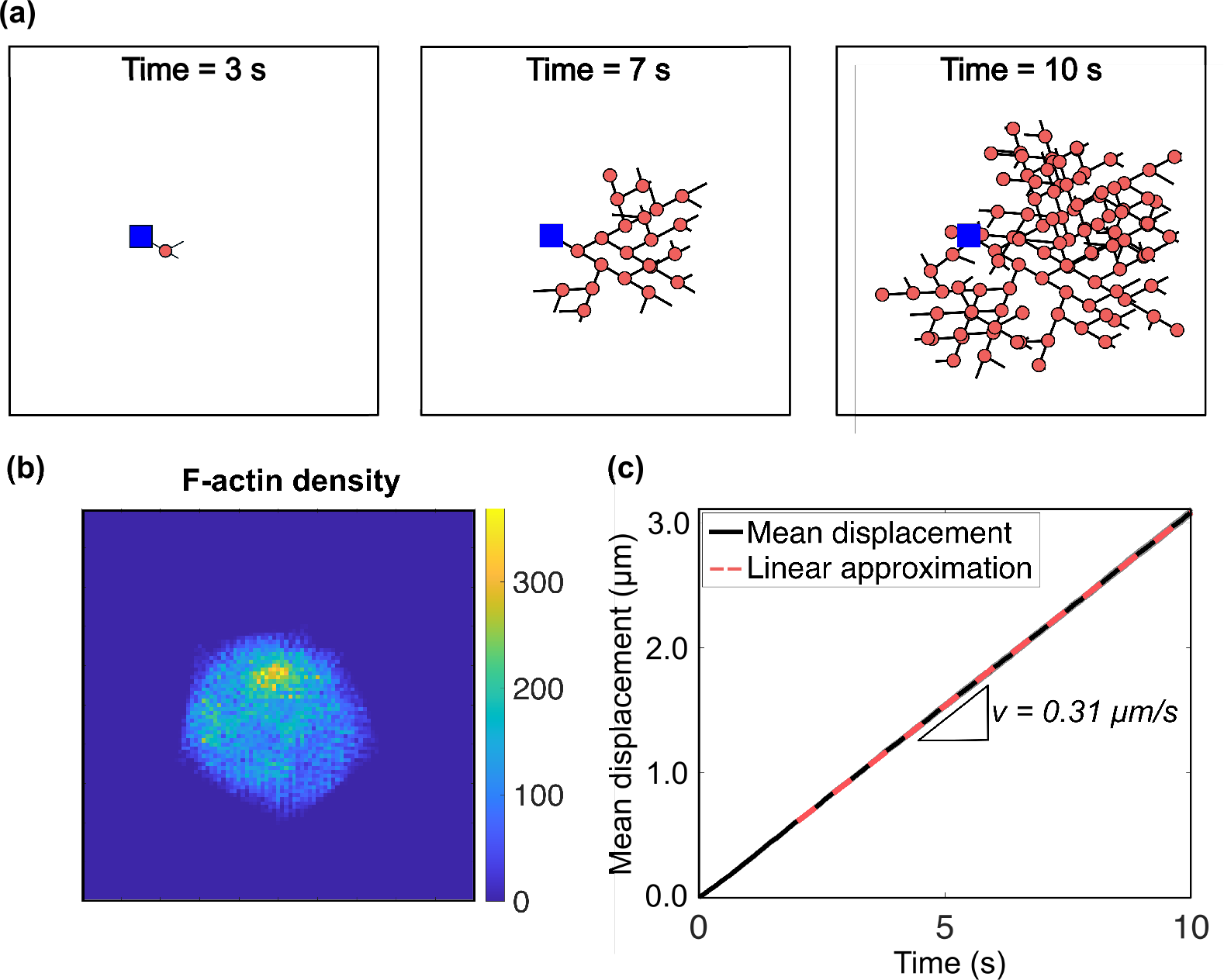
Agent-based microscale model of a branching actin structure. (a) Resulting branching networks at different time instances of *t* = 3, 7, and 10 seconds. Red dots represent Arp2/3 protein complexes, blue squares indicate the initial nucleation site, and solid black lines denote F-actin filaments. (b) F-actin length density at *t* = 10 seconds. The filament length density is calculated from one realization of the stochastic system and measures the filament length per area. (c) Mean displacement of 10 independent realizations of the model (solid black), with the corresponding best-fit linear approximation (dashed red line). The slope of the linear approximation corresponds to the wave speed of the leading edge of the network, *v* = 0.31*µ*m/s. The parameters used in stochastic agent-based simulations are provided in Table 1.

The wave-like behavior of an actin network has attracted considerable interest in recent years [47–50]. In [50], a variety of experimental and theoretical studies of actin traveling waves have been classified and reviewed. It is generally thought that actin waves result from the interplay between “activators” and “inhibitors” of actin dynamics modulated by regulatory proteins. Activation and inhibition are incorporated into our stochastic model by introducing the probabilities of branching, polymerization, and depolymerization. For a fixed set of parameter values, we can infer the values of *r*, *D* from the F-actin density averaged over many runs of the stochastic model. In addition, by varying these stochastic model parameters, we can gain insight into how they affect *r*, *D*, and the wave speed of the network.

## 3 Mathematical Methods

### 3.1 Measures to Connect Microscale Agent-Based and Macroscale Deterministic Models

In this section, we state the framework we developed to compare and connect the microscale agent-based approach in Section 2.1 to the macroscale continuum system in Section 2.2. From many instances of the stochastic simulation, the averaged mean displacement and maximal gradient of network density are computed and used, together with the analytical solution of Skellam’s equation in Eq. 24, to extract an effective bulk diffusion coefficient and unsaturated growth rate. These two quantities in the continuum model are completely identifiable by characteristics of the microscale dynamics.

#### 3.1.1 Mean Displacement

To track the movement of the actin network in the stochastic simulations, we define a “fictitious particle” to be the filament tip extending the greatest distance from the nucleation site at the origin. The position of this fictitious particle is calculated at each time step and we report the displacement of the fictitious particle as a function of time averaged over 10 independent realizations of the microscale algorithm (Fig. 2c). Note that the fictitious particle may not correspond to the same individual filament tip in consecutive time steps. We find the mean displacement over time is well-fitted by a linear function with a goodness-of-fit coefficient of *R*^2^ = 0.99996; the linear correlation coefficient varied insignificantly with parameter variations. The slope of mean displacement of the fictitious particle in the stochastic simulations can be interpreted as the speed of propagation of the leading edge of the network. The speed of the network in the continuum approach was derived in Eq. 26. Combining these yields one link between the microscale simulations and the macroscale system:

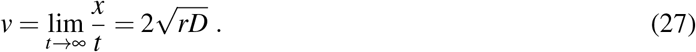

Here, *v* denotes the slope of the line which fits the mean displacement curve over time. We note that the important quantity for our analysis is “mean displacement”, not mean-squared displacement, because our system does not undergo a purely diffusive process. Instead, we use mean displacement to extract the wave speed of an advancing network that undergoes both diffusion and density-dependent growth.

#### 3.1.2 Maximal Gradient of Network Density

To distinguish between the effective diffusion coefficient and growth rate in the wave speed in Eq. 27, a second measure is necessary to isolate the two parameters. To gain insight about what the second measure should be, we look to simplify the analytic solution of Skellam’s equation to obtain an expression for one of the parameters. At an arbitrary time point *t*_*i*_, and considering a cross-section of the solution (along *y* = 0 in Fig. 2b), Eq. 24 simplifies to:

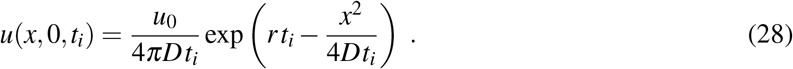

Its first and second spatial derivatives are

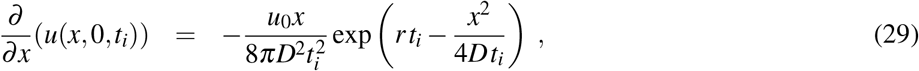

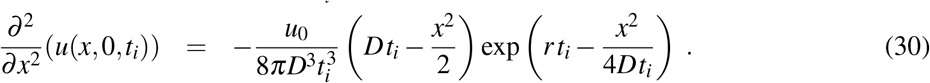

The spatial profile of the solution along a horizontal slice with *y* = 0 at *t*_*i*_ = 10 seconds together with its gradient are shown in Fig. 3a. At any point in time, there are three points of interest in the solution: the global maximum and the two inflection points. These points correspond to the zero of the gradient function (for the global maximum) and the global maxima and minima of the gradient function (for the inflection points). Critical point analysis shows that for Eq. 28 the global maximum of the solution occurs at *x* = 0, while the inflection points occur at 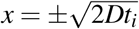.

**Figure 3:**
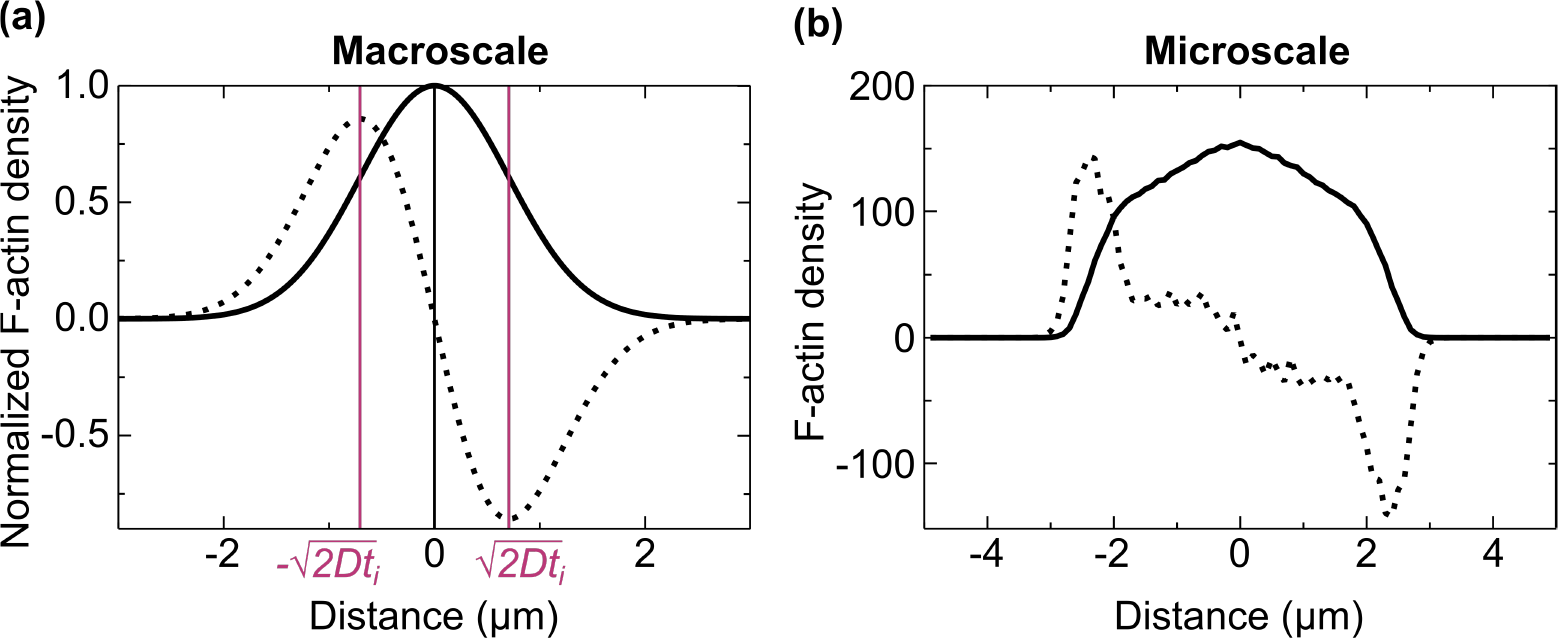
Connection between macroscale deterministic and microscale agent-based models of a branching actin network. (a) Cross-section of the solution (solid line, Eq. 28) and first derivative (dotted line, Eq. 29) of the 2D solution to Skellam’s equation with *y* = 0 at *t*_*i*_ = 10 seconds. The maximum of the solution (vertical black line) and the two inflection points (vertical magenta lines) are indicated as three points of interest used to explicitly calculate the growth rate constant (*r*) from simulations of the microscale system. (b) F-actin length density averaged over 100 independent runs of the microscale model (solid line), and its calculated gradient (dotted line).

To obtain an analytical expression for *r*, we find the global maximum of the solution curve at a time point *t*_1_:

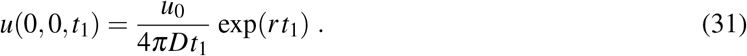

A similar expression is obtained for *t*_2_. Taking the ratio of *u*(0, 0,*t*_1_) and *u*(0, 0,*t*_2_), we conclude that:

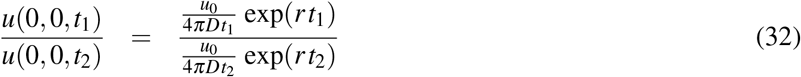

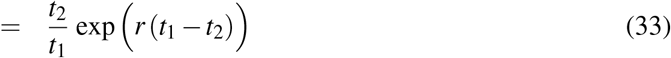

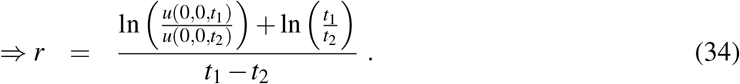

Here, *u*(0, 0, *t*_1_) and *u*(0, 0, *t*_2_) are the maximum values of our solution function at two different arbitrary time points. By averaging over many independent microscale simulations for a fixed set of parameters, we can approximate the maximum value of the actin network density. Thus, for two choices of time points *t*_1_ and *t*_2_, and the corresponding maximum values of actin network concentration averaged over many microscale simulations, we can explicitly calculate *r*, as follows:

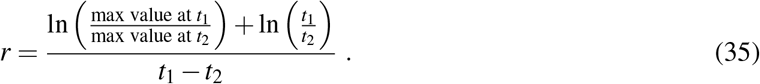

Another method for estimating *r* is to use the inflection points, rather than the maximum value of the solution profile. The inflection points at time *t*_1_ occur at 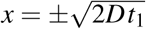. At those points, the gradient of the solution curve is:

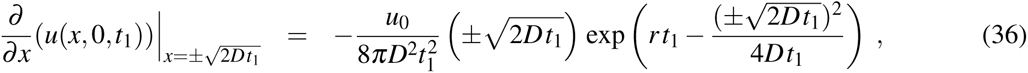

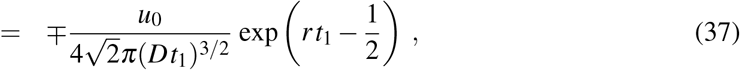

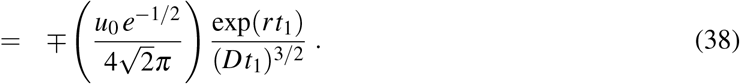

Note the first expression on the right-hand side is constant, while the second expression depends on the parameters as well as the choice of a time point. We have a similar expression at time point *t*_2_. Reminding ourselves that 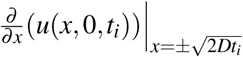 is simply the maximum gradient of the actin network at time *t*_*i*_, we take the ratio of the expressions at times *t*_1_ and *t*_2_:

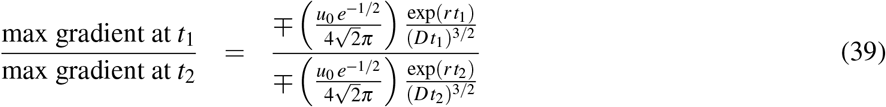

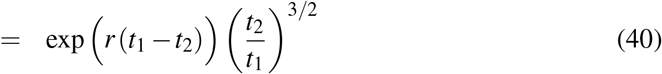

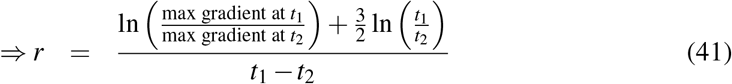

This provides a second method to explicitly calculate *r* from stochastic simulations of the microscale model.

To extract the growth rate constant *r* from the microscale agent-based system using either Eq. 35 or 41 requires a notion of actin density. To extrapolate an F-actin density from a discrete network, the computational domain is uniformly subdivided into 100 *×* 100 boxes of size 0.1 *µm* × 0.1 *µm* (Fig. 2c). To ensure a uniform distribution of the actin network, 1000 independent realizations of the microscale model are considered; at the end of each run we calculate the total length of actin filaments contained in each discretized box. Since this length is a scalar multiple of the number of F-actin monomers contained in a box, we take the length divided by the area of the box, 0.01 *µ*m^2^, to be the F-actin density at the center of that box at time *t*. The solid curve in Fig. 3b represents the averaged F-actin density at (*x*, 0) and *t* = 10 seconds with default parameters provided in Table 1. To compute its gradient, represented by the dashed line in Fig. 3b, we use centered differences to compute the spatial derivative of the calculated averaged network density.

Once we obtain *r*, and using the slope of the mean displacement of the network front, we can isolate the effective diffusion coefficient from Eq. 27 as

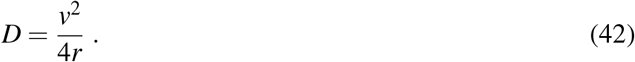

### 3.2 Sensitivity Analysis

#### 3.2.1 Numerical Implementation

To determine how microscale rates affect macroscale network behavior, we perform a series of sensitivity analyses. Specifically, we focus on three macroscopic measures: the wave speed of the advancing actin front (*v*); the effective growth rate constant (*r*); and the diffusion coefficient of the actin network (*D*) (Fig. 4). The microscale parameters varied are: the critical length required for a filament to branch (*L*_branch_); the polymerization probability (*p*_poly_), the depolymerization probability (*p*_depoly_), and the mean (*µ*) and standard deviation (*σ*) of the cumulative distribution function for filament branching probability (*p*_branch_). As several of these parameters simultaneously influence the organization of the network, three groups of parameters are established for analysis: *L*_branch_, *µ* vs. *σ*, and *p*_poly_ vs. *p*_depoly_. For each set of parameters run, all other parameters are fixed at their default values in Table 1. Parameters are varied over the following ranges: 0.15 ≤ *L*_branch_ ≤ 1.2*µm*, 0 ≤ *p*_poly_ ≤ 0.75, 0 ≤ *p*_depoly_ ≤ 1, 0 ≤ *µ* ≤ 5, and 0 ≤ *σ* ≤ 5.

**Figure 4:**
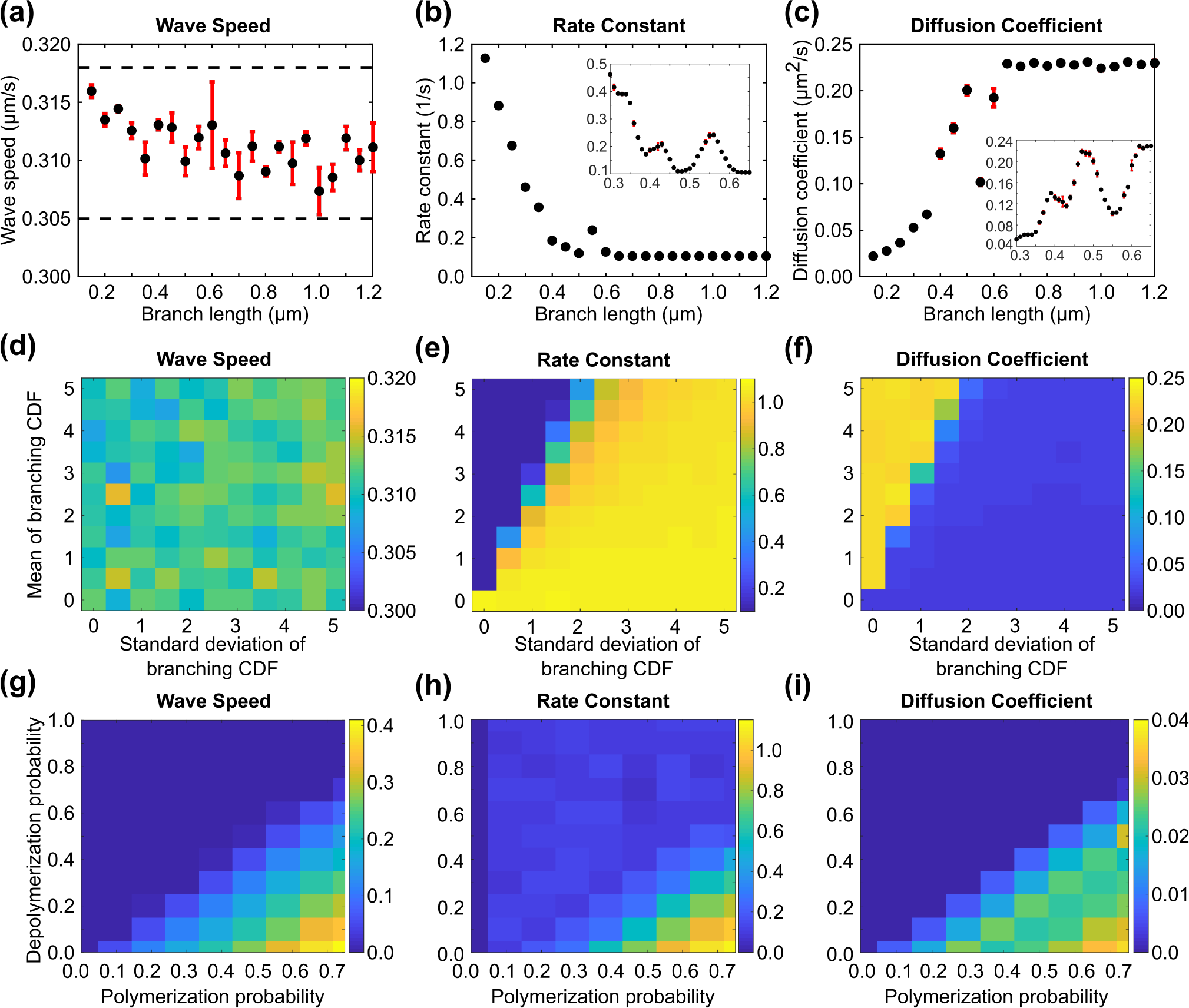
Sensitivity analysis of the wave speed (*v*, *µ*m/s), network growth rate (*r*, 1/s), and diffusion coefficient (*D*, *µ*m^2^/s) as microscale parameters are varied. Effect of critical branching length (*L*_branch_) on extracted (a) wave speed, (b) rate constant, and (c) diffusion coefficient. Black dots indicate the mean of 3 independent runs and red bars indicate standard error. The dotted lines serve as visual aids for the range of values for resulting wave speeds. The insets in (b) and (c) are more refined parameter variations for critical branching lengths between 0.3 and 0.7 *µ*m. The horizontal axes of the insets represent critical branching lengths, while the vertical axes are the growth rate constant and diffusion coefficient, respectively. Effect of parameters associated with the branching probability with mean (*µ*) and standard deviation (*σ*) on calculated (d) wave speed, (e) network growth rate, and (f) diffusion coefficient. Effect of polymerization and depolymerization probabilities, (*p*_poly_ and *p*_depoly_, respectively), on extracted (g) wave speed, (h) network growth rate, and (i) diffusion coefficient.

The lower bound for the critical branching length is chosen to ensure computational tractability – as this value is further lowered, the network density continues to grow exponentially and becomes computationally intractable. The upper bound for branching length is selected to capture the leveling-off behavior in Fig. 4b, c. Further, the interval for critical branching length encompasses many of the experimentally measured lengths. The upper bound for *p*_poly_ is again chosen to ensure computational tractability – a high polymerization rate with simultaneous low depolymerization rate increases the computational cost. Lastly, the two parameters associated with the cumulative distribution function for branching probability are non-negative. Their upper bound is arbitrarily, yet importantly captures the essential trends in the network behavior in Fig. 4e, f.

## 4 Results

### 4.1 Micro-to-Macroscale Connection

To connect the dynamics of a branched actin network across the distinct scales, we simulate 1000 runs of the microscale agent-based model and record the resulting actin density over a computational domain [−5, 5] × [−5, 5] at time points *t* = 7, 8, 9, 10, 15 seconds. We then calculate the wave speed, followed by the network growth rate and effective diffusion coefficient based on the data collected from these simulations. Specifically, the wave speed results are calculated from the average speed of a fictitious particle at the leading edge of the network, or the slope of the mean displacement over time (Fig. 4a, d, g). To obtain the slope, the plot of the mean displacement over time is fitted by a line with a goodness-of-fit coefficient of *R*^2^ = 0.9999 − 0.99999 for most choices of parameters. The goodness-of-fit is lower, *R*^2^ = 0.5 − 0.6, for similar rates of polymerization and depolymerization. This is because the network undergoes periods of growth followed by decay and the effect is even more dramatic when depolymerization rate is faster than polymerization rate. In this case we report that the wave speed is zero since overall the network cannot grow.

The maximum averaged filament length density or the maximum rate of change of the averaged density at time points *t*_1_ = 9 and *t*_2_ = 10 yields the growth rate constant from Eq. 35 (Fig. 4b, e, h). Lastly, the effective diffusion of the bulk network can be readily calculated according to Eq. 42 (Fig. 4c, f, i). We simulate the PDE model in Eq. 6 using the diffusion coefficient and growth rate determined above, and compare the density predicted by the continuum model to the averaged density produced by the agent-based model (Fig. 5).

**Figure 5:**
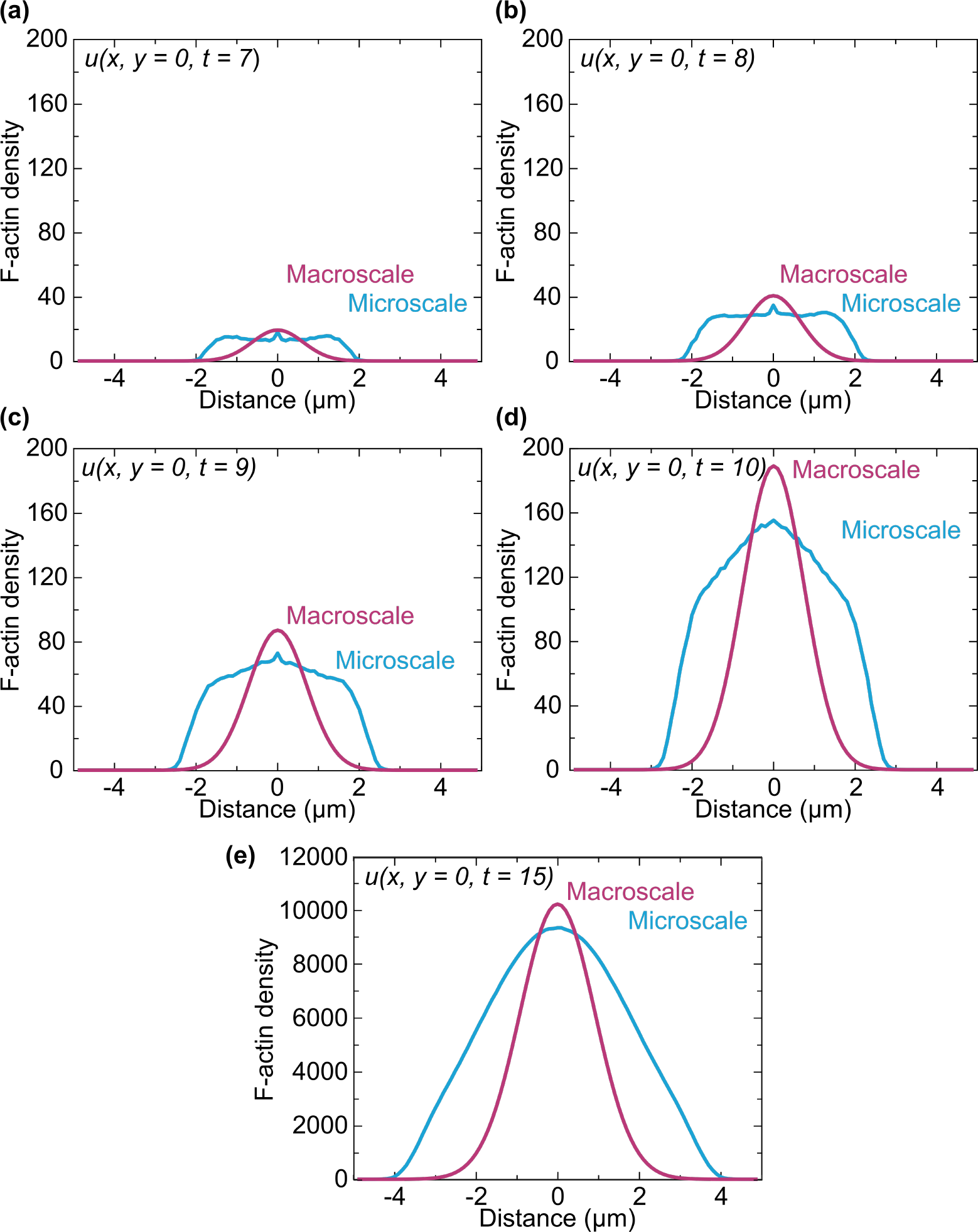
Comparison of F-actin density profiles obtained from agent-based simulations (blue) and analytic solution to Skellam’s equation (magenta) along a cross-section with *y* = 0 and time instances of (a) 7 seconds, (b) 8 seconds, (c) 9 seconds, (d) 10 seconds, and (e) 15 seconds. Macroscale parameters for the PDE, *D* = 0.03 *µm*^2^/s and *r* = 0.8 1/s, were calculated from averaged agent-based simulations as described in Section 3.

We report that the wave speed is approximately 0.31 *µ*m/s, *r* ≈ 0.8 1/s using using either Eq. 34 or Eq. 41, and *D* ≈ 0.03 *µ*m^2^/s. The averaged actin densities along a cross-section with *y* = 0 at times *t* = 7, 8, 9, 10, 15 seconds produced by the agent-based simulations are plotted in blue in Fig. 5, while the corresponding densities from Skellam’s equation are shown in magenta.

### 4.2 Results of the Sensitivity Analysis

We find that the wave speed of the actin network is largely unaffected by branching parameters, either the critical branching length or the mean and standard deviation of the cumulative distribution function for branching probability (Fig. 4a, d). However, the wave speed does depend on polymerization and depolymerization probabilities, which dictate the rate of growth of filaments. This result is reasonable given our initial assumption of unlimited resources – branching events control the spatial distribution of the network, while the rates of filament length change dictate the speed of network extension. Thus, increasing the polymerization probability increases the rate at which the network grows outwardly, i.e., the wave speed (Fig. 4g). Similarly, as the depolymerization probability increases, actin filaments are less likely to grow until eventually the overall growth of the network is arrested (top left corner on Fig. 4g).

In contrast, the effective growth rate constant and diffusion coefficient are affected by all five microscale kinetic parameters while keeping the wave speed of the network constant (Fig. 4, last two columns). We find that the growth rate constant is approximately inversely proportional to the critical branching length (Fig. 4b). The physical intuition is that the growth rate constant is a measure of the number of tips available for growth events (Eq. 7). For large critical branching lengths, the network is composed of a small number of filaments that persistently grow until the length condition for a branching event is met. Since a branching event can only occur at tips of filaments in our model, for large critical lengths only a small number of tips are available as branching sites, resulting in a small number of sites undergoing growth. The growth rate ranges between roughly 0.1 and 1.2 1/s, where the lower bound is attained for critical branching lengths over ~ 0.5 *µ*m. Variation of the two parameters of the branching cumulative distribution function – mean and standard deviation – produce a transition from the lower to the upper bound of growth rate (Fig. 4e). For a fixed but low standard deviation of the cumulative distribution function, decreasing the mean shifts the cumulative distribution to the left and thus increases the probability that a branching event can occur. However, for a fixed mean, increasing the standard deviation of the cumulative distribution decreases its slope, and thus results in larger probability for a filament to branch. Taken together, a left-shifted, shallow branching cumulative distribution function results in more branching events, and thus more filament tips that can undergo growth (bottom right in Fig. 4e). A similar but more gradual transition is found with changes in polymerization and depolymerization rates (Fig. 4h). The upper bound of the effective growth rate occurs for rapidly growing actin filaments, where polymerization probability is high but depolymerization probability is low. (Fig. 4h, bottom right). This is due to filaments with a high polymerization probability reaching their critical lengths more quickly, allowing branching to start sooner, and the network to spread out more quickly. To summarize, our findings indicate that the growth rate of the leading edge of the network is dependent on the growth and decay rates of the filaments, but also on the number of filament tips available for binding of G-actin monomers.

The effective diffusion constant is calculated from the wave speed and growth rate constant using Eq. 42. Thus, to maintain a constant wave speed as critical branching length and branching probability are varied (Fig. 4a, d), the parameter dependence of the diffusion constant must complement the parameter dependence of the rate constant (Fig. 4b-c, e-f). We find that the diffusion coefficient increases with increasing critical branching length until it reaches a plateau value of approximately 0.25 *µ*m^2^/s for branching lengths over ~ 0.5 *µ*m (Fig. 4c). In this regime, the network topology is composed of few, long filaments that grow persistently since branching does not occur until a large critical length is reached. The network front moves in an approximately ballistic way rather than a diffusive, space-exploring way. We note that the transition to a plateau occurs at a similar critical branching length of 0.5 *µ*m for both the diffusion constant and growth rate constant because the wave speed is constant at this critical branching length (in fact, it is constant across all branching length values). A sharp transition in the diffusion constant is reported as the branching probability parameters – mean and standard deviation – are varied (Fig. 4f). A high mean coupled with a low standard deviation results in a cumulative distribution function that is steep and shifted to the right. This results in a lower probability to branch, which means that individual filaments grow more persistently, and the effective diffusion of the actin network from the nucleation site is faster (top left in Fig. 4f). Reducing the mean or increasing the standard deviation increases the probability to branch, which results in a denser actin network that does not diffuse as far from the nucleation site (see smaller diffusion coefficients in bottom and right of Fig. 4f). For fixed branching parameters, the effective diffusion can be slightly increased through faster growth of the filament, or slightly decreased with faster decay of the filament length (Fig. 4i). Specifically, permissible diffusion constants ranges between 0.005 and 0.035 *µ*m^2^/s.

## 5 Discussion

Distinct profiles arise from the stochastic, microscale simulations and continuum, macroscale model (Fig. 5). In the microscale approach, the flat filamentous actin profile with sharp shoulders at the boundary indicates that the outward drive of the advancing actin network dominates over the filament production term. In contrast, the continuum model reveals a more balanced outward diffusion with reaction production at the origin, as evidenced by the smooth profile growing in both radial extent and magnitude. This functional mismatch between the microscale and macroscale results could be due to assumptions of either model. In the microscale model, we have only incorporated polymerization and depolymerization dynamics along with Arp2/3-mediated branching, while neglecting molecular motors and regulatory proteins involved in actin dynamics. Moreover, we have neglected the effects of capping of barbed ends and mechanical properties of actin filaments. Future extensions of the model will include incorporating a wider set of proteins acting on the actin filaments, as well as other physical properties such as limited availability of G-actin monomers. The latter case of constraining the cytoplasmic monomer pool presents an interesting resource-limited case study relevant for various biological conditions that will be analyzed in future work. These modifications may provide a closer match to the continuum model. With the macroscale model, we have arrived at the simplified reaction-diffusion equation in Eq. 6 by assuming both a slow actin depolymerization rate and an unlimited monomer pool. In the future, we would like to consider relaxing these conditions on the system in the hope of an improved match between the two scale models. Finally, a spatially-dependent reaction term may be incorporated to correct for the fact that the polymerization/depolymerization and especially branching reactions are not truly homogeneous reactions occurring throughout the bulk phase of the system, but rather, there is a distinct location dependence as to where the reaction is taking place (e.g., only at the filament tips or with a minimum spacing). Taken together, these results suggest that great care must be taken to ensure models of actin dynamics are consistent with the underlying physical system.

## Notes

#### Summary of Updates

Edited contribution of authors

